# Density-Yield Relationship of an Invasive Annual Grass in California Chaparral and Its Restoration Implications

**DOI:** 10.64898/2026.09.21.751057

**Authors:** Zhenyu Li, Stephanie Ma Lucero, Carla D’Antonio

**Author notes:** Massachusetts Institute of Technology, Cambridge, Massachusetts, USA. Corresponding Author Zhenyu Li. Open Research Statement All data and code used for the manuscript (Li et al., 2026) are available on Zenodo: https://doi.org/10.5281/zenodo.22286989.

## Abstract

The density-yield relationship of wild plant populations, which in agricultural systems is typically consistent with the constant final yield hypothesis, has remained largely unexplored in a restoration context despite its potential importance. In degraded California shrublands, non-native annual grasses are associated with reduced native shrub recruitment. By studying their density-yield relationship, we can potentially optimize current restoration practices by identifying the level of grass control needed to reduce the competitive pressure they create for woody plants. This in turn could enhance woody plant establishment. Here, we conducted field experiments at a degraded chaparral site in Santa Barbara County, California. We manipulated the density of *Bromus diandrus*, a dominant invasive annual grass, through 0-100% removal treatments to test its density-yield relationship. Simultaneously, we outplanted *Ceanothus megacarpus* seedlings as phytometers while also monitoring soil moisture and light availability throughout the growing season. Our results demonstrate that the *Bromus diandrus* density-yield relationship follows the constant final yield hypothesis, with strong intraspecific competition observed from moderate to high grass density contributing to a constant population biomass and a density dependence allometric exponent comparable to previous agricultural studies. A reduction in grass biomass (yield) was observed at 75% grass removal, yet an increase in resource availability (i.e., light and soil moisture) was only observed at 100% grass removal, as indicated by both direct measurements and enhanced phytometer performance. Applied to restoration in California Chaparral and other Mediterranean ecosystems, where the reduction of “weed” biomass is a common management goal, our findings suggest full removal of non-native annual grasses is likely needed to aid in the restoration of native woody seedlings.

## Introductions

The density-yield relationship has been an important consideration in plant population biology for many decades (Harper, 1977; Weiner & Freckleton, 2010), and it is important for understanding and predicting how plants will respond individually and in communities to factors that influence their survival, growth and competitive interactions. One such factor is the physical manipulation of plant density during invasive species control efforts in restoration or land management. Invasive species control efforts are typically aimed at reducing, rather than eradicating, an unwanted species and are often conducted to create temporal windows of opportunity for native species to capture freed-up resources and successfully establish into a site (Neil & Ariml, 2011; Sher et al., 2018). Yet densely growing invasive species may follow a constant final yield relationship (Weiner & Freckleton, 2010), whereby the removal of some of the invaders allows the remaining individuals to compensate for the removed biomass by increasing their own biomass, thereby limiting freed-up resources. Similarly, recovery of an invader after initial complete removal could follow a relationship whereby low-density individuals achieve high per capita biomass, with total plot biomass being equal between low- and high-density recovery plots (Shinozaki & Kira, 1954; Postma et al., 2021). This hypothesis has been repeatedly demonstrated in agronomy, with species such as buckwheat (*Fagopyrum esculentum*) and maize (*Zea mays*) amongst many others identified to follow a power law relationship and exhibiting constant final yield above a particular density (Johnson et al., 1983; Li et al., 2013; Friedman, 2016; Cao et al., 2024; Deng et al., 2006). While the concept of constant final yield has implications for invasive species management and control, it has largely been tested in agricultural settings and rarely in resource-limited natural systems (White & Harper, 1970; Xue & Hagihara, 1999). To our knowledge, it has not been applied to the context of invasive plant control and restoration.

While being an iconic part of the central and southern California’s natural landscape and key to the region’s ecosystem functions (Jennings, 2018; Lambert et al, 2010; Safford et al, 2018; Keely, 2018; Rundel, 2018), the chaparral ecosystem in California has been heavily degraded in recent years due to a combination of factors such as urbanization, short-interval fire, drought and alien annual grass invasions, giving rise to the need for restoration (Keeley & Brennan, 2012; Syphard et al., 2019; Keeley, 2001). European annual grasses, introduced in the early days of European settlement of western North America, have become widely invasive across arid and semi-arid California ecosystems (D’Antonio & Vitousek, 1992; Baker & Halsey, 2022). With traits associated with fast growth, such as high specific leaf area, high relative growth rate, and rapid seed production, they have been demonstrated to suppress recruitment of native species across California (e.g., Heady, 1977; Graebner et al., 2012; Sandel & Dangremond, 2012; Wright et al., 2004). Further, fueled by increased fire frequency, more severe drought stress, and anthropogenic activities that create ignition and disturbances (Baker & Halsey, 2022; Dewees et al., 2022; Lucero et al., 2021), these non-native annual grasses have contributed to the type-conversion and degradation of many valuable and diverse native California ecosystems (Park et al., 2018; Barro & Conard, 1991; Pratt, 2022), including the chaparral.

Concerns over the degradation of chaparral (Syphard et al, 2018) have led to efforts to revegetate or restore it (Dewees et al, in press; Allen et al., 2018; Stylinski and Allen, 1999). Yet, control or at least temporary reductions of non-native grasses are required to promote recovery of woody plants in these systems (Stratton, 2004; VinZant, 2019), as the grasses have been shown to suppress the germination and decrease the survival of native species through competition for space and water, among other valuable resources, playing a key role in limiting restoration success (Phillips & Allen, 2024; Funk et al., 2016; Engel et al., 2019). However, the extent of non-native grass removal needed to facilitate restoration success is currently unclear. By testing the application of the constant final yield hypothesis to invasive grass growth or specifically to non-native annual grasses in degraded chaparral ecosystems, we can potentially improve the efficiency of restoration by understanding what level of grass control is needed to enhance woody plant establishment.

In this study, we tested whether invasive grasses in a degraded foothill shrubland environment exhibited behavior consistent with the constant final yield hypothesis, at what density constant final yield was achieved, and how grass density and subsequent biomass interacted with the outplanting of seedlings of a once-dominant native shrub species. We quantified the density-yield relationship of a common invasive annual grass species, *Bromus diandrus,* in a type-converted non-native grass-dominated site by directly manipulating grass density, monitoring relevant environmental variables, and using outplanted *Ceanothus megacarpus*, a common chaparral species, as a phytometer for a native woody species response to possible freed-up resources. We aimed to evaluate the density at which non-native biomass is reduced, if any, and whether such a biomass reduction can lead to greater native woody seedling performance. We address the following questions: (1) What is the density-yield relationship of non-native annual grasses, to what degree does it follow the constant final yield hypothesis, and is the relationship consistent across years? (2) How does non-native annual grass thinning influence environmental resources such as light and water that could be available to other species? and (3) Can out-planted native seedlings benefit from grass reduction below full (100%) removal? Invasive grass control often involves mowing, grazing, or herbicide use, each of which reduces abundance or biomass, but from which grasses typically recover. Thus, our manual density reduction followed by regrowth is not unlike the windows of opportunity that management actions create in these systems.

## Methods

### Study Site

We conducted our study at the San Marcos Foothills Preserve in Santa Barbara, California (34°24’50’’ N and 119°50’44’’ W, elevation: 125-156 meters; Figure 1a). Research took place in 2024 (year 1) and 2025 (year 2) at two distinct sites within the Preserve (Figure 1b). Both sites occurred on south-facing slopes that were historically co-dominated by coastal sage scrub and chaparral plant communities. At the time of our study, however, they had been type-converted to non-native annual grasses and were dominated by nearly monospecific stands of *Bromus diandrus*. *Avena sp.* and *Brassica nigra* were also present but estimated to make up less than 5% of the canopy cover at each site. The site is in a Mediterranean climate with a mean annual precipitation of 546 mm and a standard error of 37 mm in the past 60 years (County of Santa Barbara). In our experimental years, total precipitation of 693 mm occurred during the 2024 growing season between December 2023 and May 2024, notably above the annual average for the location, and 374 mm during the 2025 growing season between December 2024 and April 2025 (Funk et al. 2015; Appendix S1: Figure S3). Most recently, the site was burned in December of 2019 by the Cave Fire.

**Figure 1.**
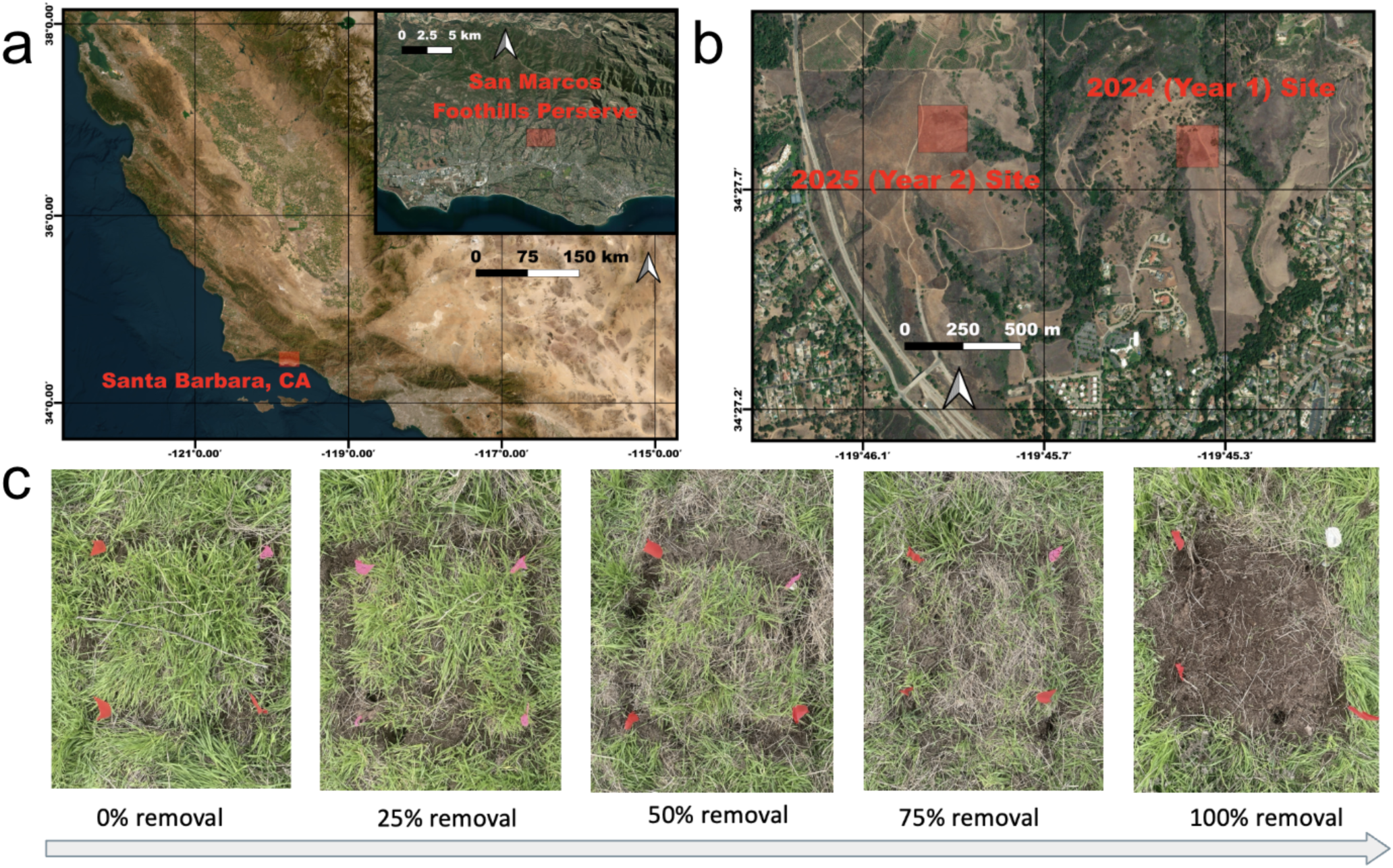
Map of the location of the field site in San Marcos Foothills Preserve (a) and map of the specific location where year 1 (2024) and year 2 (2025) of the density manipulation experiment took place (b). Photos of 50 x 50cm plots of Bromus diandrus density gradient established after administering different levels of removal treatment (c).

Soil types at both sites were located on terraces with moderately well-drained soil and were derived from mixed alluvium parent material. Soils ranged from fine sandy loam near the soil surface to clay deeper within the soil profile. However, soils at the 2025 site had greater stone and rock inclusions. Soils at the 2024 site were Milpitas-Positas fine sandy loam soils (Positas and similar soils 48%, Milpitas and similar soils 45%) while soils at the 2025 site were Milpitas stoney fine sandy loam (Milpitas and similar soils 85%) (Web Soil Survey 2026).

### Non-native Removal Treatment

On February 3rd, 2024, we set up 25 of the 50 x 50 cm plots in the 2024 site with the precise location marked on the map (Figure 1b), which had no observable shrub recruitment and was covered by newly germinated *Bromus diandrus* with no other species visible following January rainfall (Appendix S1: Figure S4). Initial non-native grass density was quantified by counting the number of grass seedlings in the marked plots. Five different treatments-0% removal, 25% removal, 50% removal, 75% removal, and 100% removal by our grass seedling count (n = 5 per treatment = 25 plots) were randomly assigned to the plots and calculated based on each plot’s original density to eventually establish a density gradient (Figure 1c). The post-removal density was multiplied by 4 to be reported as plants/m^2^ for the rest of the study. After the initial setup, the plots were allowed to grow naturally until their seeds were harvested in late April 2024, and the rest of the biomass was harvested on May 21st, 2024, when the grasses were visibly at the end of their lifecycle. We initially predicted biomass allocation would differ between reproductive and vegetative biomass. We found, however, that grass density had no effect on biomass allocation, as the reproductive and vegetative biomasses correlated strongly (R^2^ = 0.920), so the two biomass values were combined for downstream analysis in 2024 and not repeated in 2025. The combined biomass samples were dried at 60 degrees Celsius for at least 48 hours to obtain the total dried biomass for each sample.

On February 16th, 2025, we set up 50, 50 x 50 cm plots in a separate site but with identical characteristics such as *Bromus* coverage, soil type, and management history as the 2024 site (Figure 1b). Initial density survey and treatment methods were consistent with year 1 using a random design to establish a similar density gradient but with more replicates (n = 10). One, 4-month-old *Ceanothus megacarpus* seedling was randomly selected from our UCSB initiated plants (see below), and hand-planted into the center of each plot as a phytometer and watered with 400 ml of tap water. The grasses removed during the phytometer out-planting were taken into account and subtracted from the plot’s final density count, and then the systems were allowed to grow naturally. On May 30th, 2025, all of the invasive grass biomass and the phytometers were harvested separately, and both were dried in the lab at 60 degrees Celsius for at least 48 hours to obtain dry biomass.

### Soil Moisture & Light Measurements

Environmental data were tracked in each plot throughout the growing season during the second year of the experiment, from mid-February to early May 2025. For soil moisture, a time domain reflectometer (TDR model 6050X3K1, MiniTrase Kit; SoilMoisture Equipment Corp.; Santa Barbara, CA) was used to measure soil moisture using 15-centimeter probes. Surveys were performed 4 times across the growing season (February 22, March 4 and 27, and May 5th).

Three readings were taken at 3 random locations in each plot and averaged for each plot. The percent of photosynthetically active radiation that passed through the grass canopy (%PAR) was measured with a Line Quantum Sensor (LI-250A Light Meter; LI-COR Biosciences; Lincoln, NE), where the overhead meters were placed above the canopy to measure background, and three point measurements, randomly sampled, were taken below the canopy on the soil surface in each plot on each sampling date. %PAR was calculated by dividing the below-canopy point measurement PAR average by the above-canopy background PAR, to account for canopy conditions as grasses grew to peak biomass. Surveys were conducted on February 25th and March 8th and 31st, each day at around 11 AM on clear-sky days. Note that not all plots were surveyed on each date due to difficulty in locating our flags, used to mark the location of plots, when the grasses were too tall. Data monitoring stopped after March 21st because no further changes in canopy cover were observed in our field site.

### Phytometer outplanting

*Ceanothus megacarpus* was chosen as a ‘restoration’ species as it is present in remnant populations at the site today and is a typically seeding species threatened by grass invasion of former shrublands (Montygierd-Loyba & Keeley, 1987). Seeds of this shrub were collected in the San Marcos Foothills Preserve by Channel Island Restoration. Seeds were soaked in boiling water and then treated with cold-moist stratification for 30 days. After germination, seeds were transplanted into individual pots with standard potting soil mix (1 Mycorrhizae Growing Mix [PRO-MIX Gardening; Quakertown, PA]: 2 perlites: 2 Cactus Potting Mix [Miracle-Gro; Marysville, Ohio]) on October 28th, 2024. Subsequently, the seedlings were kept outdoors at the UCSB Biology Greenhouse and watered regularly without fertilizer until they were outplanted to the field on February 16th, 2025, when they were roughly 4.5 months old and without secondary growth.

To assess seedling condition after out-planting, we surveyed their stomatal conductance using a fluorometer (LI-600 Porometer/Fluorometer; LI-COR Biosciences; Lincoln, NE). Measurements were performed by clamping the leaf to fully cover the chamber and taking automatic readings. The fluorometer was zeroed regularly in the air. Data were collected on March 4th, 8th, 21st, and May 5th, and 30th, 2025. The stomatal conductance (g_sw_) value was measured and recorded for each phytometer that was still alive. Some phytometer stomatal conductance recordings were missed due to phytometer mortality, the phytometer being alive but lacking a leaf that could completely cover the measuring chamber, or our inability to locate the plot.

### Statistical Analysis

All statistical analyses were performed in R (version 4.6.0) with the tidyverse package (Wickham, 2019) and illustrated with ggplot2 (Wickham, 2016).

To characterize the relationship between invasive grass density (N) and aboveground dry grass biomass (DMa), common density-yield models were fitted to each year’s data. First, the power law model (DMa = aN^b^) was calculated by the method described in previous literature (Friedman, 2024), through linearizing the relationship via log-log transformation (log(DMa) = log(a) + b·log(N)) and applying ordinary least squares regression, where a and b are fitted coefficients. Based on the power law model result of the 2024 data, a linear model (DMa = aN + b) was fitted to only the 2024 data using simple regression. Based on the power law model result for 2025 data, an asymptotic model (DMa = DMa_max · (1 − exp(−2.257 · (N / N̅)))) was also fitted to only the 2025 data, where DMa_max is the observed maximum biomass and N̅ is the mean density (Friedman, 2024). Model fit for the power law and linear model was evaluated using the coefficient of determination (R²).

To calculate an overall trend for non-native annual grasses, as none of the models calculate a threshold between grass biomass and density, biomass data from both years were combined, and an estimated threshold was calculated based on the average biomass of the top fifteen (20%) samples (by biomass) from both years’ data combined.

Removal treatments (0%, 25%, 50%, 75%, 100%) were used as a categorical variable to evaluate the effect of grass removal on the measured response variables, including soil moisture, %PAR, stomatal conductance, and phytometer biomass. One-way analysis of variance (ANOVA) was conducted for each sampling date. Normality of each response variable within each treatment group was assessed using the Shapiro-Wilk test. A high proportion of %PAR data groups violated the requirements for normality, likely due to the bounded nature of the variable (constrained between 0 and 1). The arcsine square root transformation did not improve normality. As a result, Kruskal-Wallis tests were run in parallel with ANOVA for all PAR sampling dates, and conclusions were compared to reveal largely similar results, so ANOVA results were retained for consistency. For all data types, when ANOVA indicated a statistically significant treatment effect (p < 0.05), pairwise differences between treatment levels were assessed using the Tukey-Kramer honest significant difference (HSD) post-hoc test, which accounts for the familywise error rate across all pairwise comparisons.

To assess whether invasive grass density was a significant predictor of environmental conditions on individual sampling dates, grass density count post-treatment was used as a continuous, independent variable, and simple linear regressions were fit between it and each of the following response variables: soil moisture, light availability (%PAR), and stomatal conductance on all sampling dates. Regressions were run on two subsets of data points: all plots combined and all plots excluding the 100% removal treatment, to evaluate whether the presence of fully cleared plots disproportionately influenced the relationship. The significance of each regression was evaluated using the F-test p-value, and goodness of fit was reported as R².

## Results

### Density-yield relationship of non-native annual grass

Density of the grasses varied dramatically between the two years of the study, with maximum density in 2024 (year 1) being 548 plants/m^2^ (Figure 2a) while in 2025 (year 2) density was as high as 2856 plants/m^2^ (Figure 2b). The two years also differed in the shape of the density-yield relationship. Power law model on the 2024 grass biomass data produced an allometric exponent of 0.956 (1 indicates linearity), which signaled a lack of constant final yield within the range of densities for that year (Figure 2a). With the linear model, we found that aboveground dry grass biomass was strongly correlated with non-native grass density (R^2^ = 0.713). The 2025 biomass data, by contrast, exhibited an expected constant yield relationship with an exponent of 0.636. The decreased exponent signals a stronger leveling-off effect at high density, and such a relationship was also fitted to the asymptotic model (Figure 2b).

**Figure 2.**
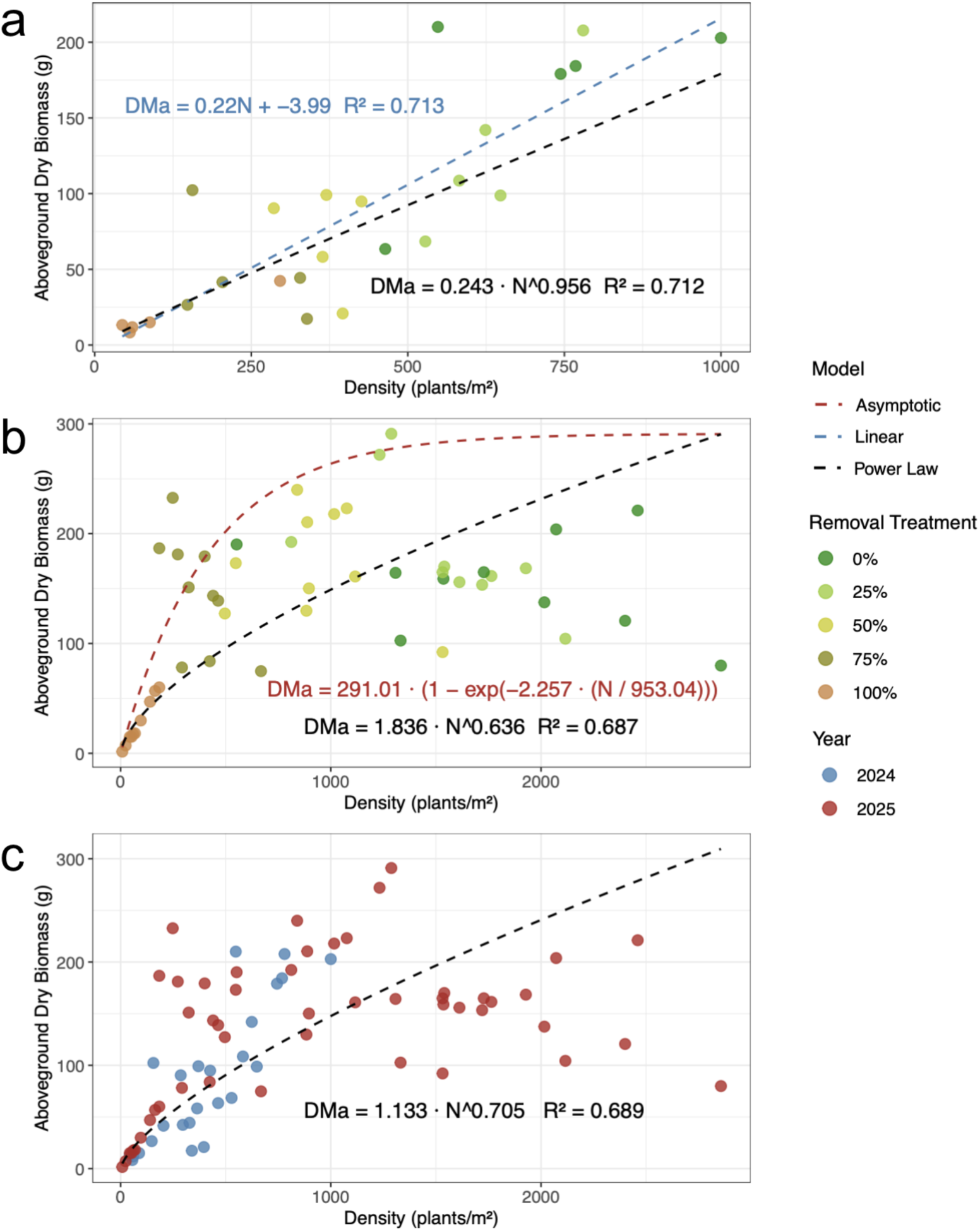
Density-yield relationship of non-native annual grass *Bromus diandrus* with data from 2024 (a), 2025 (b), and both years’ data overlaid together (c). The y-axis indicates the total aboveground dry biomass (grams), with both vegetative and reproductive growth, for each plot harvest at the end of its growing season. The initial thinning treatment producing the density is indicated by colors in panels a and b, with 0% being no removal and 100% being full removal. Panel c shows 2024 (blue) and 2025 (red) data overlaid on top of each other, with colors indicating the year. The three models applied, linear (blue line), asymptotic (red line), and power law (black lines), are shown alongside the data points, and the parameters that produced the corresponding dotted lines are shown in the equations of the same colors.

The combined data showed similar carrying capacity and overall density-yield relationship for the grass population despite previously described differences, as the biomass was visibly similar in the density range where they overlapped (Figure 2c). The calculated biomass threshold for constant yield is 220 +/- 29 grams, which was generally reached at around 500 plants/m^2^ which corresponds with approximately 75% removal treatment. The threshold can be reached as early as 184 grass seedlings per square meter (see data point with 75% removal that produced 186.67 grams of biomass). After reaching the apparent population biomass threshold, the subsequent increase in population density did not result in an increase in biomass. The population biomass remained around the biomass threshold as the density reached around 2500 grasses per square meter, but higher density plots began to show a decrease in biomass (Figure 2c) with plots at the high end of density producing values such as 79.89 and 120.67 grams, much lower than the threshold biomass.

### Density effects on resources

The abiotic environment strongly responded to our grass density manipulations. %PAR monitoring demonstrated low light penetration unless the *Bromus* population density was reduced by the 100% removal treatment (Figure 3b). In the early growing season on February 25th and March 8th, significant differences were found between the 100% removal treatment and all other treatments on February 25th (Appendix S1: Table S1). On March 8th, the 100% removal treatment remained significantly higher than the rest of the treatments (Appendix S1: Table S1), and the 75% removal treatment was also significantly higher compared to the 0% removal treatment (p = 2.52e-3). Towards the middle of their growing season, on March 21st, similarly low %PAR was found in 0% to 75% removal plots, signifying dense cover and intense shading, significantly lower than the %PAR of 100% removal treatment plots (Figure 3b; Appendix S1: Table S1).

**Figure 3.**
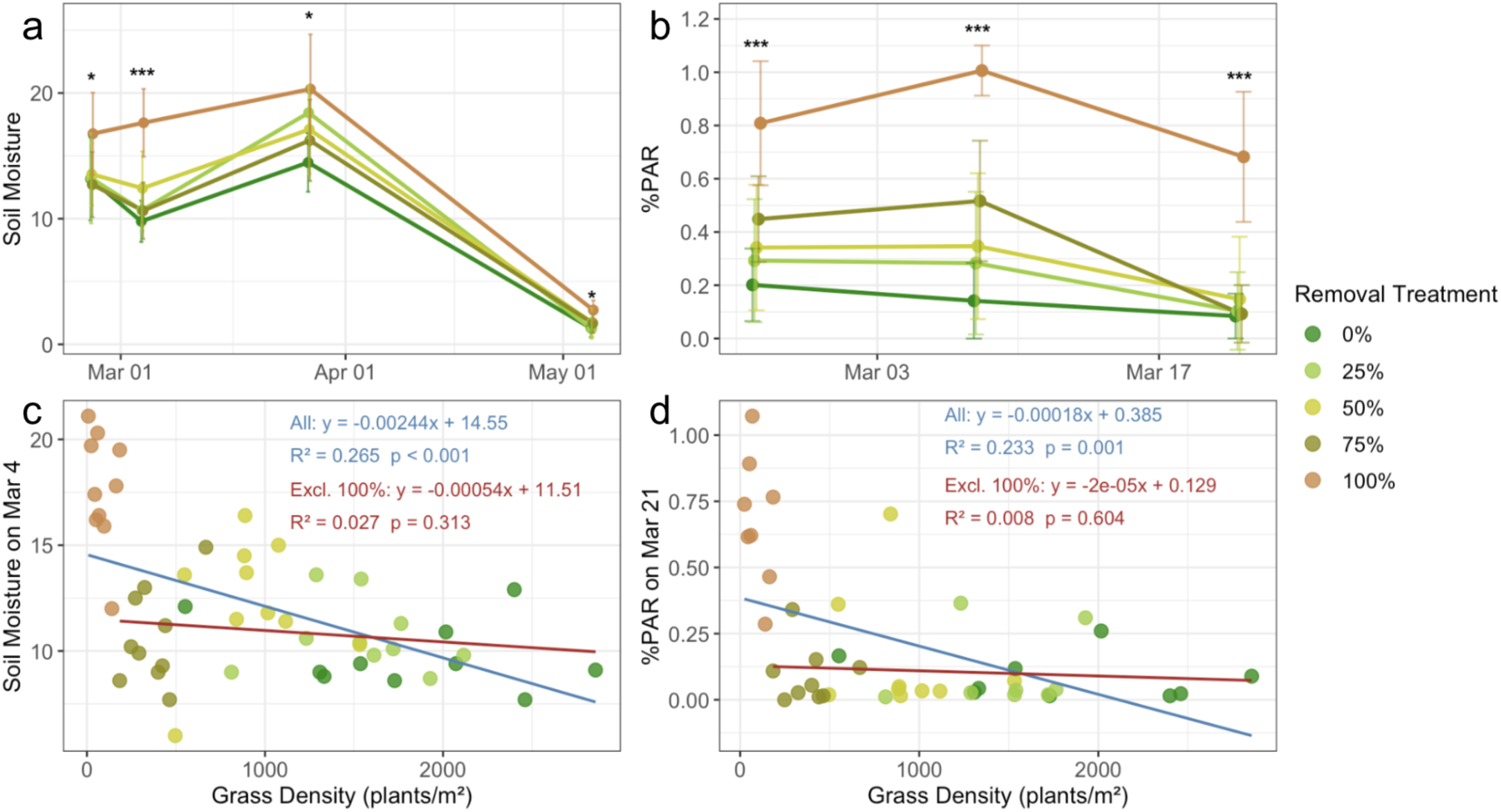
2025 field environmental data monitoring measures the effect of removal treatment on (a) soil moisture and (b) PAR through the *Bromus diandru*s growing season. Data separated by initial removal treatment, indicated by color. The mean for each treatment at each date is indicated by a point, and the error bar represents plus or minus 1 standard deviation. The result of ANOVA for each date is noted on the figure, with * = p < 0.05 ** = p < 0.01 *** = p < 0.001. Continuous density effect on soil moisture (c) and %PAR (d) at a single date, March 4th for soil moisture and March 21st for % PAR. Linear regression analysis and the statistical results are embedded in the figure, with blue showing the relationship between environmental data and grass density for all data points, and red showing that of only the 0% to 75% removal treatment.

Soil moisture also only consistently increased in 100% removal plots. Data on February 25th showed a significant difference between 100% and 75% removal only (p = 0.039). Subsequent March 4th survey showed much stronger differences between 100% and all the rest of the removal treatments (Appendix S1: Table S1). The March 27th survey also showed significant differences but to a weaker effect, with only a significant difference between 0% and 100% removal found (p = 0.012). In the May 5th survey, soil moisture was much drier compared to prior surveys as summer approached, but 100% removal was still significantly higher than the 0% (p = 0.016) and 25% removal (p = 0.021) treatments (Figure 3a).

Abiotic data on each date collected were also plotted against continuous grass density (post treatment) instead of discrete treatments. Linear regression of March 4th data showed a significant negative association between soil moisture and grass density, but no significant relationship was detected when the 100% removal plots were excluded (Figure 3c). A similar trend was observed in the %PAR data collected on March 21st (Figure 3d), as full data showed a significant negative correlation that suggests light level increases as grass density decreases, but no such relationship was found when only looking at the 0% to 75% removal data. Albeit with less statistical significance, the continuous density effect on abiotic environmental data showed a visually similar trend in other dates as well (Appendix S1: Figure S1).

### Phytometer growth and physiological response

To measure the response of native species to grass removal treatments, the removal also resulted in the end-of-season dried aboveground phytometer biomass being greater in the 100% removal plots compared to each of the phytometers in the 0% to 75% removal treatment plots (Figure 4a). The mean biomass of phytometer seedlings grown in plots with extremely low density produced by 100% removal was 0.426 grams ± 0.249, with maximum biomass being 0.848 grams. Phytometer biomass in the 100% removal plots was significantly greater than the rest (p ≤ 0.047).

**Figure 4.**
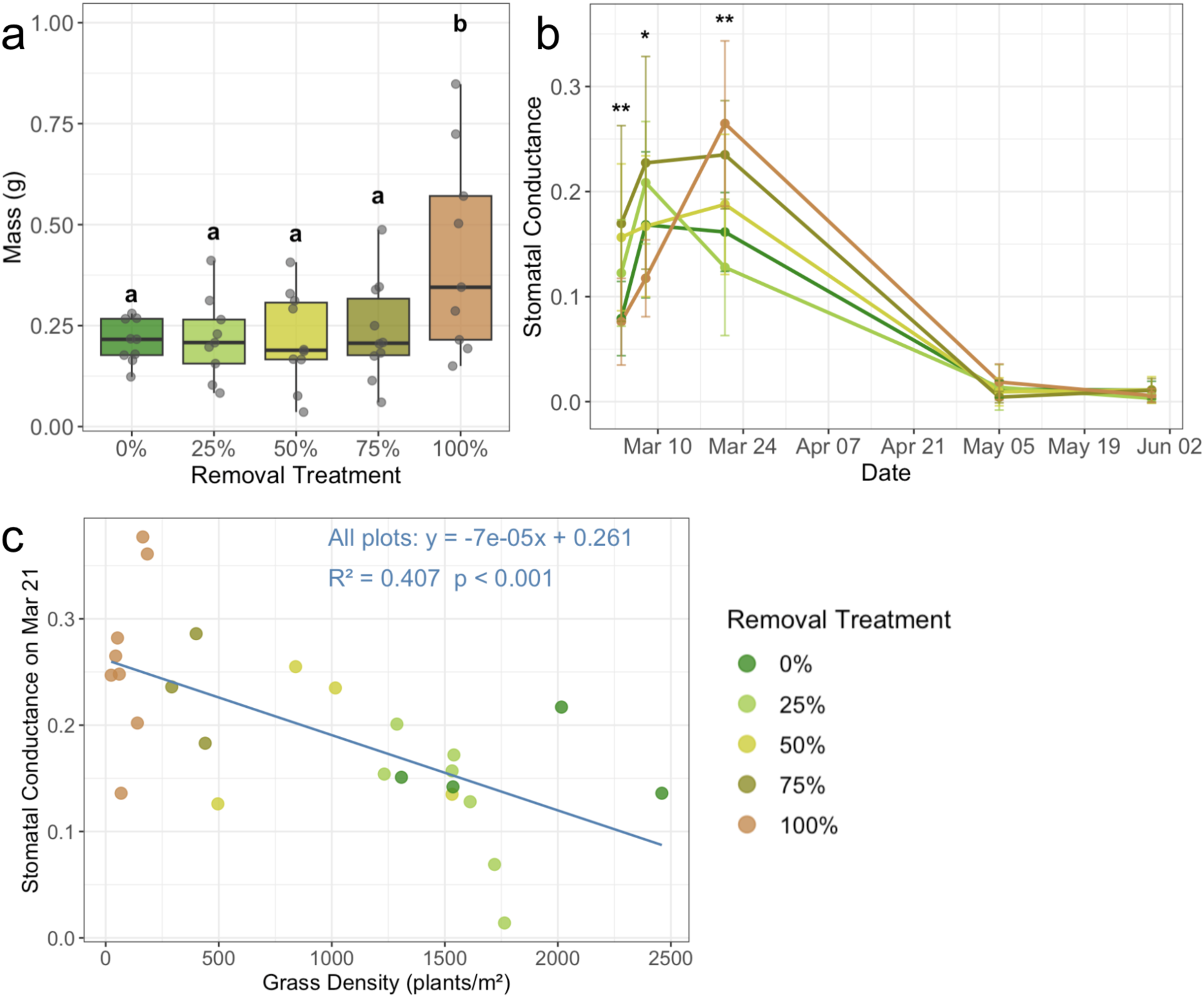
Biomass for outplanted native shrub seedlings of *Ceanothus megacarpus* at the end of the season (a). Data separated by initial removal treatment indicated by color, and letters indicate significant differences with p < 0.05. Stomatal conductance (g_sw_) value measured for each phytometer throughout the grass growing season is also shown (b). The mean for each treatment at each date is indicated by a point, and the error bar represents plus or minus 1 standard deviation. The result of the ANOVA for each date is noted on the figure, with * = p < 0.05 ** = p < 0.01 *** = p < 0.001. (c) Continuous non-native grass density effect on native seedling stomatal conductance at a single date, March 21st.

Phytometer stomatal conductance value measured on March 4th, 24 days after outplanting, showed surprisingly significant greater stomatal conductance between 75% and 0% removal (p = 0.017) and 75% and 100% removal (p = 0.016), and the March 8th data showed differences between 100% and 75% removal (p = 0.012). A linear regression analysis on the continuous grass density on phytometer stomatal conductance, however, found no significant correlation between the variables on these two days (Appendix S1: Figure S2). The subsequent survey on March 21st showed average phytometer stomatal conductance to be the highest in 100% removal plots (Figure 4b), significantly greater than the 25% removal treatment (p = 0.005). Linear regression showed a significant negative association between phytometer stomatal conductance and grass density (Figure 4c). End-of-season stomatal conductance data collected on May 5th and May 30th showed low average stomatal conductance and no differences across treatments (Appendix S1: Table S2).

## Discussion

### Density-yield relationship of non-native annual grass

In plots dominated by *Bromus diandrus*, a wide-spread non-native annual grass species that can have detrimental impacts in chaparral and other resource limited systems (Germino et al., 2016), our study showed an overall similar density-yield relationship between 2024 and 2025 despite differences in their maximum density. The overlapped region (≤1000 grass/m^2^) for the two years is largely linear, suggesting there were uncaptured resources by the population (Harpole et al., 2007). Such observation is consistent with the initial linear growth described by the constant final yield hypothesis (Weiner & Freckleton, 2010). After reaching the apparent population biomass threshold at high grass densities (>1000 grass/m^2^), the subsequent increase in population density did not result in an increase in population biomass due resource limitation (Mrad et al., 2000) and intraspecific competition (Friedman, 2016b; Zhai et al. 2018), as predicted by the constant final yield hypothesis. Furthermore, a decrease in population biomass was observed as density increases beyond that amounts suggesting high intraspecific competition and consistent with density-dependent mortality or self-thinning in a resource limited environment (Willey & Heath, 1969; Yahuza, 2011; Fibich et al., 2014; Assefa et al., 2018; Deng et al., 2012). Together our study supports previous studies demonstrating that a constant final yield can occur in natural, non-agricultural systems (Xue & Hagihara, 2008; Xue & Hagihara, 1999).

The visual constant final yield relationship among *B. diandrus* can be supported by the comparable calculated coefficients relative to previous studies across ecosystems. In the power law model initially proposed by Kira et al. (1953), the allometric exponent at the onset of growth is predicted to be 1, and the 0.958 (Figure 2a) calculated from data before the population reached constant final yield is in excellent agreement with the proposed value. For populations that reached the constant final yield, the mean exponent found by Postma et al. (2021) was 0.276, and Friedman (2024b) found exponents ranging from 0.086 to 0.535 in agricultural systems, lower than the value (0.636) found in our studies. The comparatively larger allometric exponent for *B. diandrus* suggests a weaker constant final yield effect compared to other species, potentially due to the lack of metabolic flexibility associated with some annual grasses (McBride et al., 2024), or it could also simply be explained by species specificity (Pretzsch, 2006). Notably, neither equation used fully captures the density-yield relationship we observed, as higher-density data points are all systematically lower than those predicted by the models, suggesting strong density-dependent forces, such as intraspecific competition, can operate in this setting.

The larger allometric exponent in our study could also be due to experimental differences in how we created our grass density gradient and at what life stage intraspecific competition took place. A field experiment allowed us to study grasses in situ, but it also meant we carried out thinning treatments among already germinated grasses as opposed to creating a density gradient through sowing grass seed at different densities (as in Li et al., 2013, 2016). This difference in creating a density gradient meant intraspecific competition took place later with young, month-old grass individuals compared to earlier with germinating grass seedlings, potentially weakening its effect.

### Density effects on resources

We focused on water and light because they are the most likely abiotic resources that limit non-native grass growth, and both %PAR and soil moisture responded to grass density manipulation.

With resources are presumably depleted in the control 0% removal treatment, the similar levels for 25%, 50%, and 75% treatments with the control on multiple dates indicate similar depletion, revealing compensatory responses by the remaining individuals (Ortega et al., 2004; Retuerto & Woodward, 2001). Such results resonate with the previously discussed constant final yield, as, for example, when the individuals in low-density plots exhibit compensatory growth to increase in size, a closed canopy is created to decrease the detected %PAR value (Champion et al., 1998; Whaley et al., 2000). Similar results are observed with soil moisture, suggesting water is depleted and also limits overall productivity in California grassland environments despite previous studies having indicated otherwise (Houlton & Field, 2010; Dukes et al., 2005).

Additionally, we found that only the 100% removal treatment is effective in increasing environmental resource availability, as the significant correlation between resources and grass density was lost when 100% removal plots were excluded, indicating that complete removal was necessary to increase resource availability in the environment substantially (Figure 3). Quickly consumed and heavily limiting across a wide range of density from 0% to 75% removal, light and water likely play key roles in mediating plant-to-plant interaction and are the center of intraspecific competition at our site, which is expected for the generally resource poor, semi-arid mediterranean climate of California chaparral land (Abraham et al, 2008; Pratt et al., 2007; Molinari & D’Antonio, 2014), eventually to play a major role in limiting grassland productivity at our site and contribute to the previously discussed constant final yield. Other studies have found nutrients, such as nitrogen and phosphorus availability, also influence grassland productivity (Borer et al., 2017; Houlton & Field, 2010; Chu et al., 1996). It is very likely that the interaction between nutrients and water may together drive the productivity of California non-native grassland, and they should be considered in future studies.

### Phytometer growth and physiological response

The altered resource availability, such as soil moisture, at different grass densities also influences interspecific competition, mediating native species establishment in degraded systems (Thaxton et al., 2011). The significant increase in *Ceanothus megacarpus* biomass only at 100% non-native grass removal echos the low resource availability found at all but 100% removal, and it reflects the expected beneficial effects of grass removal that have been demonstrated across multiple systems (Figure 4; Silletti et al., 2004; Cabin et al., 2002). More specifically, the importance of 100% removal supports Eliason & Allen (1997), where they found 4-month-old *Artemisia californica* seedling biomass only increased at 0%-5% non-native density. However, other studies have found 50% removal to be similarly effective, such as Philips & Allen (2024) with 6-month-old *Adenostoma fasciculatum* seedling and Rehm et al. (2023) with germinated *Coprosma rhynchocarpa* seedlings all benefited from both 50% and 100% non-native grass removal. Therefore, factors such as seedling size, systems’ nutrient richness, and species together influence the outcome of interspecific competition, but when implementing young seedlings in a source poor environment, like this study, 100% removal may be necessary in specific cases.

From a stomatal conductance perspective, no clear trend was observed in early-season, likely due to transplant shock before the phytometers had adapted to their environment (Howard and Minnich, 1989; Allen et al., 2019). Mid-season data (March 21st) suggested phytometers in 100% removal plots are more metabolically active than those in any other treatment similar to results of other studies (Jiang et al., 2006; Gago et al., 2016). Possibly, the extended increase of soil moisture and light availability in the 100% removal plots (Figure 3) allowed such a trend to persist in 100% removal plots between March 21st and May 5th, retaining greater stomatal conductance that may led to the eventually greater biomass (Roche, 2015; Kimura et al., 2020). Additionally, with similarly low stomatal conductance among all individuals, the end-of-season data suggest grass removal does not improve the environmental stressors commonly faced by native species such as drought (Jacobsen & Pratt, 2018), which has been shown to lower stomatal conductance in chaparral shrubs (Lunneberg et al., 2023) and is reflected in our study. Overall, stomatal conductance data provide further evidence that biologically available resources created by non-native grass removal, especially during the mid-growing season, can be captured by neighboring native species when intensely removed and improved their competitive advantage against non-natives through the increased growth that was observed.

### Limitations & Restoration implications

Our study has several limitations. First the removal treatment was conducted approximately one month after wet up, rendering the germinated grasses more established. Grass size will influence competition between the grasses and the outplanted *C. megacarpus* seedling (Thompson, 1988). We may have observed less suppression if the phytometer outplanting had occurred at an earlier grass stage. Additionally, our study only evaluated the response of one phytometer species, and functional trait differences among native shrub species could create different physiological responses towards the treatments leading to different outcomes (Ackerly, 2004; Nielsen et al., 2019, Dewees et al. in press). However, by using a common species found in chaparral communities but not often used in a research context due to challenging germination cues required (Hadley, 1961), this study adds to the existing knowledge of shrub species that would benefit from grass removal.

From a restoration perspective, our study supports a minimum 75% removal of grasses is necessary to reduce the invasive grasses population biomass. In other words, efforts to control non-native grasses that do not keep it reduced below 75% will not lead to a reduction in overall grass productivity. After initial grass removal efforts, regermination, which commonly occurs due to a generally large non-native seedbank, and continued rainy season precipitation, will require management attention to keep grass densities below 500 individuals/m^2^. Although grass biomass will decrease at 75% removal, to release the environmental resources to aid out-planted native seedling growth, 100% removal treatments may need to be implemented based on the response of *Ceanothus megacarpus*, as we found no environmental resource release or phytometer growth benefit in any other removal treatment but 100% (Figure 3; Figure 4). In the other removal treatments, it is likely that the remaining grasses, after thinning, compensate for the resources freed by thinning treatments, resulting in no benefit for the phytometer.

## Conclusions

This study provides a framework for considering the restoration of native woody plants in non-native annual grass dominated land through the lens of the constant final yield hypothesis. Our findings support the existence of a constant final yield in *Bromus diandrus* grass populations in degraded California chaparral, driven by intense intraspecific competition for resources but starting at relatively low density. Applied to restoration, where a reduction of biomass is desired to aid native seedling establishment, we found that an increase in resource availability and an increase in phytometer biomass can only be achieved when non-native annual grasses are fully removed, despite there being a reduction in non-native grass biomass at 75% removal. Full (i.e., 100%) non-native grass removal may be achieved with herbicides at the outset of the rainy season after grass germination, but our results suggest that any further grass germination could start to reduce woody seedling performance.

## Acknowledgments

The authors are grateful to the UCSB Coastal Fund (grant number CF-202310-07489 and CF-202409-10767), and to the Schuyler Endowment in Environmental studies for financial support. The authors thank Lauren Harris, John Fan, Emily Tian, Lily Vine, Andrea Estrada, Jason Han, Max Tu, Emely Valdez, Ash Cheng, Jackie Calhoun, and Amalia Narofsky for providing critical field work assistance. We also thank the D’Antonio lab group for feedback throughout the project from conception to final analyses.

## Conflict of Interest Statement

The authors declare no conflicts of interest.

## Notes

### Competing Interest Statement

The authors have declared no competing interest.

