## Appendix 1 for "Density-Yield Relationship of an Invasive Annual Grass in California Chaparral and Its Restoration Implications"

Supporting Information

ECOSPHERE
AN ESA OPEN ACCESS JOURNAL

Subject Track - Vegetation Ecology

**Manuscript Title**

Density-Yield Relationship of an Invasive Annual Grass in California Chaparral and Its Restoration Implications

**Authors**

Zhenyu Li, Stephanie Ma Lucero, Carla D'Antonio

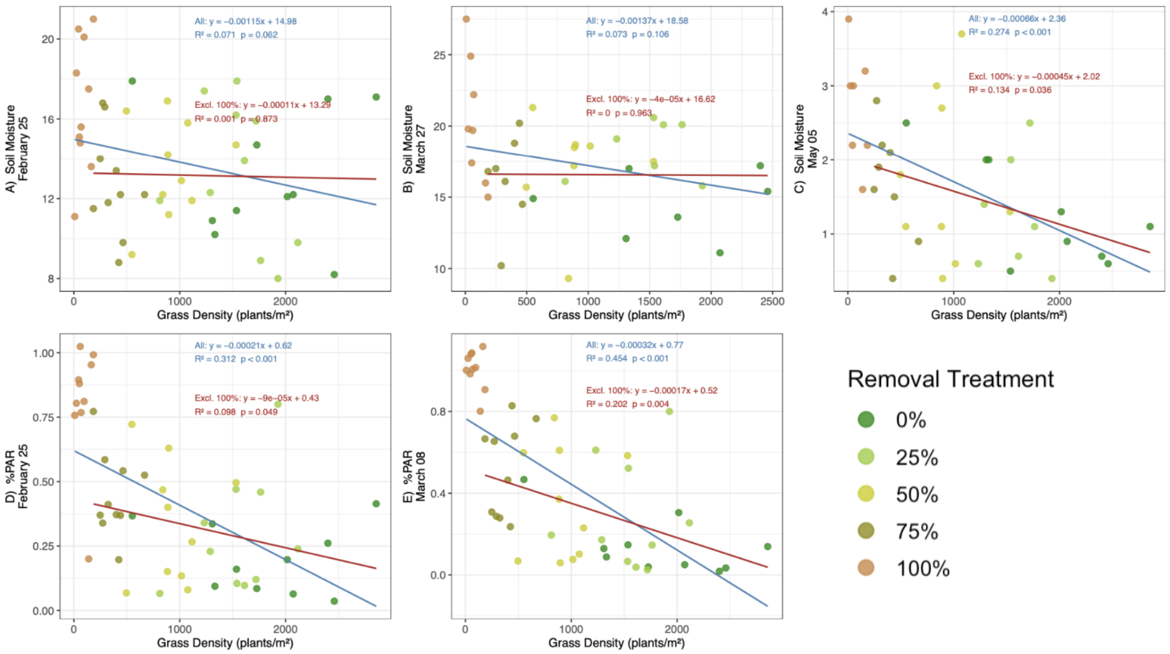


**Figure S1.** Multi-panel figure showing %PAR or soil moisture by non-native density for the dates shown in Figure 3.

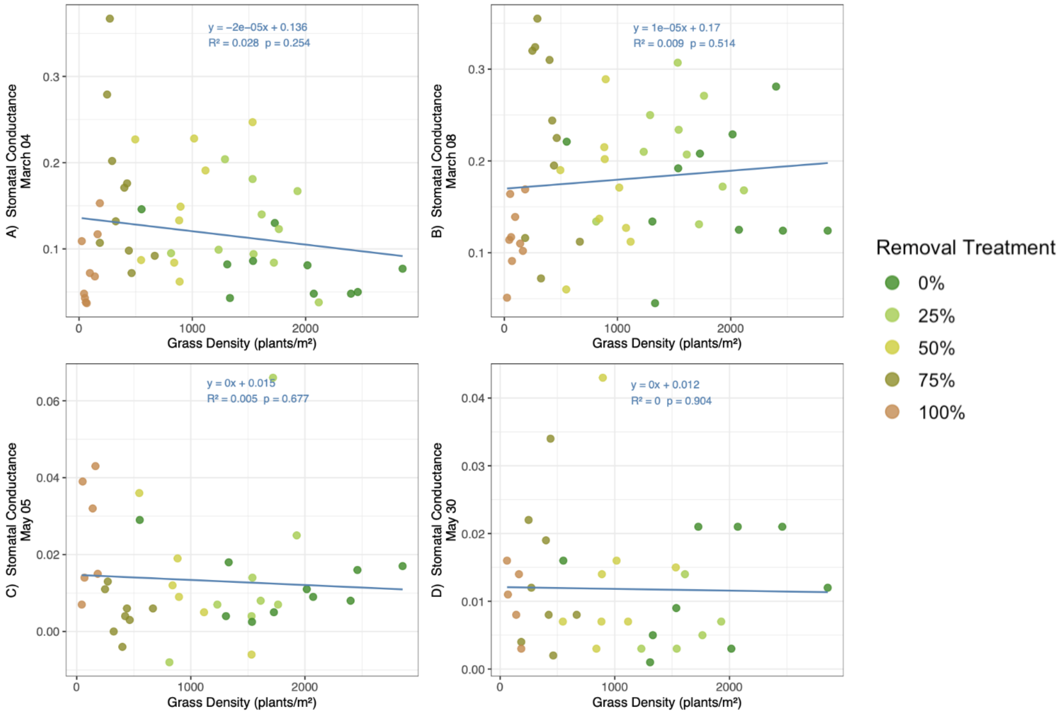


**Figure S2.** Multi-panel figure showing phytometer stomatal conductance by non-native density for the dates shown in Figure 4.

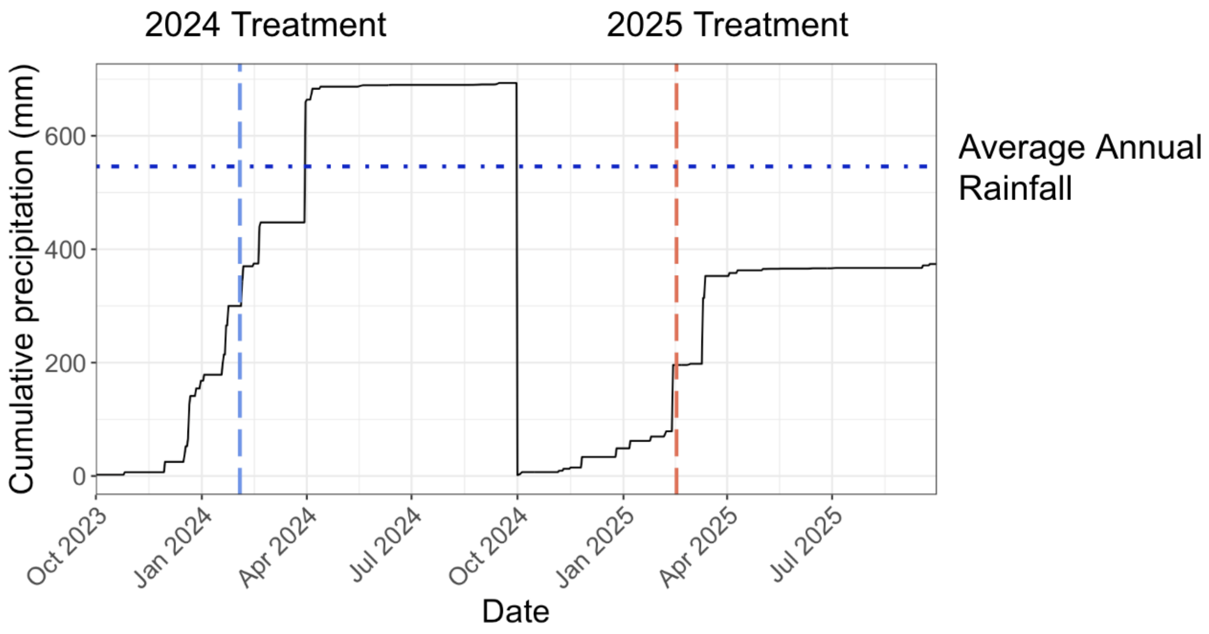


**Figure S3.** Precipitation at San Marcos Foothills Preserve (Santa Barbara) during the 2024 and 2025 growing seasons, when the experiment was conducted. Daily precipitation from 2023.10.01 - 2025.09.30 - data from CHIRPS Precipitation Daily (UCSB Climate Hazards Center InfraRed Precipitation with Station Data (Version 2.0 Final)), extracted via Google Earth Engine. Blue and red vertical lines represent dates when the treatments are implemented each year, and the blue horizontal line represents the average annual rainfall of the area.

| Variable | Date | ANOVA_p | ANOVA_sig | Comparison | diff | lwr | upr | p_adj | sig |
|---|---|---|---|---|---|---|---|---|---|
| **Light Availability (%PAR)** | Feb 25 2025 | 0 | *** | 0.25-0 | 0.0911 | -0.1678 | 0.35 | 0.8541 | ns |
| **Light Availability (%PAR)** | Feb 25 2025 | 0 | *** | 0.5-0 | 0.1401 | -0.1188 | 0.399 | 0.5442 | ns |
| **Light Availability (%PAR)** | Feb 25 2025 | 0 | *** | 0.5-0.25 | 0.049 | -0.2099 | 0.3079 | 0.9829 | ns |

| | | | | | | | | | |
|---|---|---|---|---|---|---|---|---|---|
| **Light Availability (%PAR)** | Feb 25 2025 | 0 | *** | 0.75-0 | 0.2468 | -0.0121 | 0.5057 | 0.0684 | ns |
| **Light Availability (%PAR)** | Feb 25 2025 | 0 | *** | 0.75-0.25 | 0.1557 | -0.1032 | 0.4146 | 0.4391 | ns |
| **Light Availability (%PAR)** | Feb 25 2025 | 0 | *** | 0.75-0.5 | 0.1067 | -0.1522 | 0.3656 | 0.7676 | ns |
| **Light Availability (%PAR)** | Feb 25 2025 | 0 | *** | 1-0 | 0.607 | 0.3481 | 0.8659 | 0 | *** |
| **Light Availability (%PAR)** | Feb 25 2025 | 0 | *** | 1-0.25 | 0.5159 | 0.257 | 0.7748 | 0 | *** |
| **Light Availability (%PAR)** | Feb 25 2025 | 0 | *** | 1-0.5 | 0.4669 | 0.208 | 0.7258 | 1E-04 | *** |
| **Light Availability (%PAR)** | Feb 25 2025 | 0 | *** | 1-0.75 | 0.3602 | 0.1013 | 0.6191 | 0.0024 | ** |
| **Light Availability (%PAR)** | Mar 08 2025 | 0 | *** | 0.25-0 | 0.1414 | -0.1293 | 0.4121 | 0.5778 | ns |
| **Light Availability (%PAR)** | Mar 08 2025 | 0 | *** | 0.5-0 | 0.2051 | -0.0656 | 0.4758 | 0.2164 | ns |
| **Light Availability (%PAR)** | Mar 08 2025 | 0 | *** | 0.5-0.25 | 0.0637 | -0.207 | 0.3344 | 0.9621 | ns |
| **Light Availability (%PAR)** | Mar 08 2025 | 0 | *** | 0.75-0 | 0.375 | 0.1043 | 0.6457 | 0.0025 | ** |
| **Light Availability (%PAR)** | Mar 08 2025 | 0 | *** | 0.75-0.25 | 0.2336 | -0.0371 | 0.5043 | 0.1203 | ns |
| **Light Availability (%PAR)** | Mar 08 2025 | 0 | *** | 0.75-0.5 | 0.1699 | -0.1008 | 0.4406 | 0.3957 | ns |
| **Light Availability (%PAR)** | Mar 08 2025 | 0 | *** | 1-0 | 0.8646 | 0.5939 | 1.1353 | 0 | *** |
| **Light Availability (%PAR)** | Mar 08 2025 | 0 | *** | 1-0.25 | 0.7232 | 0.4525 | 0.9939 | 0 | *** |
| **Light Availability (%PAR)** | Mar 08 2025 | 0 | *** | 1-0.5 | 0.6595 | 0.3888 | 0.9302 | 0 | *** |
| **Light Availability (%PAR)** | Mar 08 2025 | 0 | *** | 1-0.75 | 0.4896 | 0.2189 | 0.7603 | 1E-04 | *** |
| **Light Availability (%PAR)** | Mar 21 2025 | 0 | *** | 0.25-0 | 0.019 | -0.2237 | 0.2618 | 0.9994 | ns |
| **Light Availability (%PAR)** | Mar 21 2025 | 0 | *** | 0.5-0 | 0.0632 | -0.1723 | 0.2987 | 0.938 | ns |
| **Light Availability (%PAR)** | Mar 21 2025 | 0 | *** | 0.5-0.25 | 0.0442 | -0.1985 | 0.2869 | 0.9847 | ns |
| **Light Availability (%PAR)** | Mar 21 2025 | 0 | *** | 0.75-0 | 0.0082 | -0.2273 | 0.2437 | 1 | ns |

| | | | | | | | | | |
|---|---|---|---|---|---|---|---|---|---|
| **Light Availability (%PAR)** | Mar 21 2025 | 0 | *** | 0.75-0.25 | -0.0108 | -0.2535 | 0.2319 | 0.9999 | ns |
| **Light Availability (%PAR)** | Mar 21 2025 | 0 | *** | 0.75-0.5 | -0.055 | -0.2905 | 0.1805 | 0.9619 | ns |
| **Light Availability (%PAR)** | Mar 21 2025 | 0 | *** | 1-0 | 0.5979 | 0.3552 | 0.8406 | 0 | *** |
| **Light Availability (%PAR)** | Mar 21 2025 | 0 | *** | 1-0.25 | 0.5789 | 0.3291 | 0.8286 | 0 | *** |
| **Light Availability (%PAR)** | Mar 21 2025 | 0 | *** | 1-0.5 | 0.5347 | 0.292 | 0.7774 | 0 | *** |
| **Light Availability (%PAR)** | Mar 21 2025 | 0 | *** | 1-0.75 | 0.5897 | 0.347 | 0.8324 | 0 | *** |
| **Soil Moisture** | Feb 25 2025 | 0.0357 | * | 0.25-0 | 0.05 | -3.8642 | 3.9642 | 1 | ns |
| **Soil Moisture** | Feb 25 2025 | 0.0357 | * | 0.5-0 | 0.37 | -3.5442 | 4.2842 | 0.9988 | ns |
| **Soil Moisture** | Feb 25 2025 | 0.0357 | * | 0.5-0.25 | 0.32 | -3.5942 | 4.2342 | 0.9993 | ns |
| **Soil Moisture** | Feb 25 2025 | 0.0357 | * | 0.75-0 | -0.46 | -4.3742 | 3.4542 | 0.9972 | ns |
| **Soil Moisture** | Feb 25 2025 | 0.0357 | * | 0.75-0.25 | -0.51 | -4.4242 | 3.4042 | 0.9959 | ns |
| **Soil Moisture** | Feb 25 2025 | 0.0357 | * | 0.75-0.5 | -0.83 | -4.7442 | 3.0842 | 0.974 | ns |
| **Soil Moisture** | Feb 25 2025 | 0.0357 | * | 1-0 | 3.59 | -0.3242 | 7.5042 | 0.0862 | ns |
| **Soil Moisture** | Feb 25 2025 | 0.0357 | * | 1-0.25 | 3.54 | -0.3742 | 7.4542 | 0.0934 | ns |
| **Soil Moisture** | Feb 25 2025 | 0.0357 | * | 1-0.5 | 3.22 | -0.6942 | 7.1342 | 0.1521 | ns |
| **Soil Moisture** | Feb 25 2025 | 0.0357 | * | 1-0.75 | 4.05 | 0.1358 | 7.9642 | 0.0393 | * |
| **Soil Moisture** | Mar 04 2025 | 0 | *** | 0.25-0 | 0.88 | -2.0428 | 3.8028 | 0.9114 | ns |
| **Soil Moisture** | Mar 04 2025 | 0 | *** | 0.5-0 | 2.63 | -0.2928 | 5.5528 | 0.0961 | ns |
| **Soil Moisture** | Mar 04 2025 | 0 | *** | 0.5-0.25 | 1.75 | -1.1728 | 4.6728 | 0.4435 | ns |
| **Soil Moisture** | Mar 04 2025 | 0 | *** | 0.75-0 | 0.84 | -2.0828 | 3.7628 | 0.9241 | ns |
| **Soil Moisture** | Mar 04 2025 | 0 | *** | 0.75-0.25 | -0.04 | -2.9628 | 2.8828 | 1 | ns |

| | | | | | | | | | |
|---|---|---|---|---|---|---|---|---|---|
| **Soil Moisture** | Mar 04 2025 | 0 | *** | 0.75-0.5 | -1.79 | -4.7128 | 1.1328 | 0.4205 | ns |
| **Soil Moisture** | Mar 04 2025 | 0 | *** | 1-0 | 7.84 | 4.9172 | 10.7628 | 0 | *** |
| **Soil Moisture** | Mar 04 2025 | 0 | *** | 1-0.25 | 6.96 | 4.0372 | 9.8828 | 0 | *** |
| **Soil Moisture** | Mar 04 2025 | 0 | *** | 1-0.5 | 5.21 | 2.2872 | 8.1328 | 1E-04 | *** |
| **Soil Moisture** | Mar 04 2025 | 0 | *** | 1-0.75 | 7 | 4.0772 | 9.9228 | 0 | *** |
| **Soil Moisture** | Mar 27 2025 | 0.0192 | * | 0.25-0 | 3.9571 | -1.0732 | 8.9875 | 0.18 | ns |
| **Soil Moisture** | Mar 27 2025 | 0.0192 | * | 0.5-0 | 2.6286 | -2.2421 | 7.4992 | 0.5332 | ns |
| **Soil Moisture** | Mar 27 2025 | 0.0192 | * | 0.5-0.25 | -1.3286 | -6.1992 | 3.5421 | 0.9322 | ns |
| **Soil Moisture** | Mar 27 2025 | 0.0192 | * | 0.75-0 | 1.7571 | -3.2732 | 6.7875 | 0.8492 | ns |
| **Soil Moisture** | Mar 27 2025 | 0.0192 | * | 0.75-0.25 | -2.2 | -7.2304 | 2.8304 | 0.7147 | ns |
| **Soil Moisture** | Mar 27 2025 | 0.0192 | * | 0.75-0.5 | -0.8714 | -5.7421 | 3.9992 | 0.985 | ns |
| **Soil Moisture** | Mar 27 2025 | 0.0192 | * | 1-0 | 5.8411 | 0.9704 | 10.7117 | 0.0124 | * |
| **Soil Moisture** | Mar 27 2025 | 0.0192 | * | 1-0.25 | 1.8839 | -2.9867 | 6.7546 | 0.7961 | ns |
| **Soil Moisture** | Mar 27 2025 | 0.0192 | * | 1-0.5 | 3.2125 | -1.493 | 7.918 | 0.3019 | ns |
| **Soil Moisture** | Mar 27 2025 | 0.0192 | * | 1-0.75 | 4.0839 | -0.7867 | 8.9546 | 0.1352 | ns |
| **Soil Moisture** | May 05 2025 | 0.016 | * | 0.25-0 | -0.046 | -1.2899 | 1.1979 | 1 | ns |
| **Soil Moisture** | May 05 2025 | 0.016 | * | 0.5-0 | 0.4556 | -0.708 | 1.6191 | 0.792 | ns |
| **Soil Moisture** | May 05 2025 | 0.016 | * | 0.5-0.25 | 0.5016 | -0.7423 | 1.7455 | 0.7739 | ns |
| **Soil Moisture** | May 05 2025 | 0.016 | * | 0.75-0 | 0.3861 | -0.8133 | 1.5855 | 0.8851 | ns |
| **Soil Moisture** | May 05 2025 | 0.016 | * | 0.75-0.25 | 0.4321 | -0.8453 | 1.7096 | 0.8656 | ns |
| **Soil Moisture** | May 05 2025 | 0.016 | * | 0.75-0.5 | -0.0694 | -1.2688 | 1.1299 | 0.9998 | ns |

| | | | | | | | | | |
|---|---|---|---|---|---|---|---|---|---|
| **Soil Moisture** | May 05 2025 | 0.016 | * | 1-0 | 1.4397 | 0.1958 | 2.6836 | 0.0166 | * |
| **Soil Moisture** | May 05 2025 | 0.016 | * | 1-0.25 | 1.4857 | 0.1663 | 2.8051 | 0.0208 | * |
| **Soil Moisture** | May 05 2025 | 0.016 | * | 1-0.5 | 0.9841 | -0.2598 | 2.228 | 0.1773 | ns |
| **Soil Moisture** | May 05 2025 | 0.016 | * | 1-0.75 | 1.0536 | -0.2239 | 2.3311 | 0.1474 | ns |

**Table S1**. ANOVA and Tukey-Kramer test results statistics for pairwise comparison of environmental data collected in each plot by initial removal treatments. Both PAR and soil moisture comparisons are included.

| Variable | Date | ANOVA_p | ANOVA_sig | Comparison | diff | lwr | upr | p_adj | sig |
|---|---|---|---|---|---|---|---|---|---|
| **Stomatal Conductance (gsw)** | Mar 04 2025 | 0.0035 | ** | 100%-0% | -0.003 | -0.084 | 0.0781 | 1 | ns |
| **Stomatal Conductance (gsw)** | Mar 04 2025 | 0.0035 | ** | 100%-25% | -0.0464 | -0.1274 | 0.0347 | 0.4873 | ns |
| **Stomatal Conductance (gsw)** | Mar 04 2025 | 0.0035 | ** | 100%-50% | -0.0803 | -0.1635 | 0.0028 | 0.0627 | ns |
| **Stomatal Conductance (gsw)** | Mar 04 2025 | 0.0035 | ** | 100%-75% | -0.0935 | -0.1745 | -0.0124 | 0.0165 | * |
| **Stomatal Conductance (gsw)** | Mar 04 2025 | 0.0035 | ** | 25%-0% | 0.0434 | -0.0355 | 0.1223 | 0.5264 | ns |
| **Stomatal Conductance (gsw)** | Mar 04 2025 | 0.0035 | ** | 50%-0% | 0.0773 | -0.0037 | 0.1584 | 0.0678 | ns |
| **Stomatal Conductance (gsw)** | Mar 04 2025 | 0.0035 | ** | 50%-25% | 0.0339 | -0.0471 | 0.115 | 0.7556 | ns |
| **Stomatal Conductance (gsw)** | Mar 04 2025 | 0.0035 | ** | 75%-0% | 0.0905 | 0.0116 | 0.1694 | 0.0173 | * |
| **Stomatal Conductance (gsw)** | Mar 04 2025 | 0.0035 | ** | 75%-25% | 0.0471 | -0.0318 | 0.126 | 0.4447 | ns |
| **Stomatal Conductance (gsw)** | Mar 04 2025 | 0.0035 | ** | 75%-50% | 0.0132 | -0.0679 | 0.0942 | 0.9903 | ns |
| **Stomatal Conductance (gsw)** | Mar 08 2025 | 0.0155 | * | 100%-0% | -0.0509 | -0.1429 | 0.0412 | 0.5222 | ns |

| | | | | | | | | | |
|---|---|---|---|---|---|---|---|---|---|
| **Stomatal Conductance (gsw)** | Mar 08 2025 | 0.0155 | * | 100%-25% | -0.091 | -0.183 | 0.0011 | 0.0541 | ns |
| **Stomatal Conductance (gsw)** | Mar 08 2025 | 0.0155 | * | 100%-50% | -0.0496 | -0.144 | 0.0449 | 0.5718 | ns |
| **Stomatal Conductance (gsw)** | Mar 08 2025 | 0.0155 | * | 100%-75% | -0.1099 | -0.2019 | -0.0178 | 0.0122 | * |
| **Stomatal Conductance (gsw)** | Mar 08 2025 | 0.0155 | * | 25%-0% | 0.0401 | -0.0495 | 0.1297 | 0.708 | ns |
| **Stomatal Conductance (gsw)** | Mar 08 2025 | 0.0155 | * | 50%-0% | -0.0013 | -0.0933 | 0.0907 | 1 | ns |
| **Stomatal Conductance (gsw)** | Mar 08 2025 | 0.0155 | * | 50%-25% | -0.0414 | -0.1334 | 0.0506 | 0.7042 | ns |
| **Stomatal Conductance (gsw)** | Mar 08 2025 | 0.0155 | * | 75%-0% | 0.059 | -0.0306 | 0.1486 | 0.3459 | ns |
| **Stomatal Conductance (gsw)** | Mar 08 2025 | 0.0155 | * | 75%-25% | 0.0189 | -0.0707 | 0.1085 | 0.9742 | ns |
| **Stomatal Conductance (gsw)** | Mar 08 2025 | 0.0155 | * | 75%-50% | 0.0603 | -0.0317 | 0.1523 | 0.3511 | ns |
| **Stomatal Conductance (gsw)** | Mar 21 2025 | 0.0081 | ** | 100%-0% | 0.1033 | -0.0171 | 0.2236 | 0.1159 | ns |
| **Stomatal Conductance (gsw)** | Mar 21 2025 | 0.0081 | ** | 100%-25% | 0.1369 | 0.0352 | 0.2386 | 0.0051 | ** |
| **Stomatal Conductance (gsw)** | Mar 21 2025 | 0.0081 | ** | 100%-50% | 0.077 | -0.0434 | 0.1974 | 0.3448 | ns |
| **Stomatal Conductance (gsw)** | Mar 21 2025 | 0.0081 | ** | 100%-75% | 0.0298 | -0.1033 | 0.1628 | 0.9615 | ns |
| **Stomatal Conductance (gsw)** | Mar 21 2025 | 0.0081 | ** | 25%-0% | -0.0336 | -0.1568 | 0.0895 | 0.9235 | ns |
| **Stomatal Conductance (gsw)** | Mar 21 2025 | 0.0081 | ** | 50%-0% | 0.0262 | -0.1127 | 0.1652 | 0.9791 | ns |
| **Stomatal Conductance (gsw)** | Mar 21 2025 | 0.0081 | ** | 50%-25% | 0.0599 | -0.0633 | 0.1831 | 0.605 | ns |
| **Stomatal Conductance (gsw)** | Mar 21 2025 | 0.0081 | ** | 75%-0% | 0.0735 | -0.0766 | 0.2236 | 0.5988 | ns |
| **Stomatal Conductance (gsw)** | Mar 21 2025 | 0.0081 | ** | 75%-25% | 0.1071 | -0.0285 | 0.2428 | 0.1677 | ns |
| **Stomatal Conductance (gsw)** | Mar 21 2025 | 0.0081 | ** | 75%-50% | 0.0472 | -0.1029 | 0.1974 | 0.8789 | ns |
| **Stomatal Conductance (gsw)** | May 05 2025 | 0.1432 | | 100%-0% | 0.013 | -0.0078 | 0.0339 | 0.3869 | ns |
| **Stomatal Conductance (gsw)** | May 05 2025 | 0.1432 | | 100%-25% | 0.0096 | -0.0122 | 0.0314 | 0.7084 | ns |

| | | | | | | | | | |
|---|---|---|---|---|---|---|---|---|---|
| **Stomatal Conductance (gsw)** | May 05 2025 | 0.1432 | | 100%-50% | 0.0125 | -0.0108 | 0.0358 | 0.5399 | ns |
| **Stomatal Conductance (gsw)** | May 05 2025 | 0.1432 | | 100%-75% | 0.0201 | -0.0017 | 0.0419 | 0.0816 | ns |
| **Stomatal Conductance (gsw)** | May 05 2025 | 0.1432 | | 25%-0% | 0.0034 | -0.0157 | 0.0226 | 0.9851 | ns |
| **Stomatal Conductance (gsw)** | May 05 2025 | 0.1432 | | 50%-0% | 6E-04 | -0.0203 | 0.0214 | 1 | ns |
| **Stomatal Conductance (gsw)** | May 05 2025 | 0.1432 | | 50%-25% | -0.0029 | -0.0247 | 0.0189 | 0.9953 | ns |
| **Stomatal Conductance (gsw)** | May 05 2025 | 0.1432 | | 75%-0% | -0.0071 | -0.0262 | 0.0121 | 0.8221 | ns |
| **Stomatal Conductance (gsw)** | May 05 2025 | 0.1432 | | 75%-25% | -0.0105 | -0.0307 | 0.0097 | 0.5688 | ns |
| **Stomatal Conductance (gsw)** | May 05 2025 | 0.1432 | | 75%-50% | -0.0076 | -0.0294 | 0.0142 | 0.8492 | ns |
| **Stomatal Conductance (gsw)** | May 30 2025 | 0.643 | | 100%-0% | -0.0017 | -0.0169 | 0.0134 | 0.9974 | ns |
| **Stomatal Conductance (gsw)** | May 30 2025 | 0.643 | | 100%-25% | 0.004 | -0.0132 | 0.0212 | 0.9602 | ns |
| **Stomatal Conductance (gsw)** | May 30 2025 | 0.643 | | 100%-50% | -0.0036 | -0.0191 | 0.0119 | 0.9604 | ns |
| **Stomatal Conductance (gsw)** | May 30 2025 | 0.643 | | 100%-75% | -0.0032 | -0.0187 | 0.0123 | 0.9733 | ns |
| **Stomatal Conductance (gsw)** | May 30 2025 | 0.643 | | 25%-0% | -0.0057 | -0.0209 | 0.0094 | 0.8083 | ns |
| **Stomatal Conductance (gsw)** | May 30 2025 | 0.643 | | 50%-0% | 0.0019 | -0.0113 | 0.0151 | 0.9934 | ns |
| **Stomatal Conductance (gsw)** | May 30 2025 | 0.643 | | 50%-25% | 0.0076 | -0.0079 | 0.0231 | 0.6175 | ns |
| **Stomatal Conductance (gsw)** | May 30 2025 | 0.643 | | 75%-0% | 0.0015 | -0.0117 | 0.0147 | 0.9972 | ns |
| **Stomatal Conductance (gsw)** | May 30 2025 | 0.643 | | 75%-25% | 0.0072 | -0.0083 | 0.0227 | 0.6607 | ns |
| **Stomatal Conductance (gsw)** | May 30 2025 | 0.643 | | 75%-50% | -4E-04 | -0.014 | 0.0132 | 1 | ns |

**Table S2.** ANOVA and Tukey-Kramer test results statistics for pairwise comparison of phytometer stomatal conductance data collected in each phytometer plant by initial removal treatments.
